# Non-uniform contextual interactions in the visual cortex place fundamental limits on spatial vision

**DOI:** 10.1101/2023.08.15.553380

**Authors:** Mitchell P. Morton, Sachira Denagamage, Nyomi V. Hudson, Anirvan S. Nandy

**Affiliations:** Department of Neuroscience, Yale University, New Haven, CT 06510; Department of Psychology, Yale University, New Haven, CT 06511; Interdepartmental Neuroscience Program, Yale University, New Haven, CT 06510; Wu Tsai Institute, Yale University, New Haven, CT 06511; Kavli Institute for Neuroscience, Yale University, New Haven, CT 06511

## Abstract

A prevailing assumption in our understanding of how neurons in the primary visual cortex (V1) integrate contextual information is that such processes are spatially uniform. Conversely, perceptual phenomena such as visual crowding, the impaired ability to accurately recognize a target stimulus among distractors, suggest that interactions among stimuli are distinctly non-uniform. Prior studies have shown flankers at specific spatial geometries exert differential effects on target perception. To resolve this discrepancy, we investigated how flanker geometry impacted the representation of a target stimulus in the laminar microcircuits of V1. Our study reveals flanker location differentially impairs stimulus representation in excitatory neurons in the superficial and input layers of V1 by tuned suppression and untuned facilitation of orientation responses. Mechanistically, this effect can be explained by asymmetrical spatial kernels in a normalization model of cortical activity. Strikingly, these non-uniform modulations of neural representation mirror perceptual anisotropies. These results establish the non-uniform spatial integration of information in the earliest stages of cortical processing as a fundamental limitation of spatial vision.

## INTRODUCTION

Natural vision activates neurons not in isolation, but in the context of other stimuli. The extensive literature on contextual interactions in the visual cortex, particularly surround modulation (*1–9*), has significantly enriched our understanding of visual processing. However, a prevailing assumption in this literature has been the uniform nature of the surrounding context. This is evident in the experimental design of contextual stimuli typically as annuli or as a circular constellation of flankers (*1, 2, 5, 10*). In contrast, perceptual non-uniformities in spatial vision have garnered considerable interest in the past several decades (*11–14*). In particular, the polar angle with respect to the center of gaze has been shown to be especially relevant to perceptual performance (see review by (*15*)). Perceptual non-uniformities are especially striking in visual crowding, a ubiquitous phenomenon that profoundly limits the ability to recognize target objects among visual clutter (*16*). Crowding thus offers a unique window to investigate the neural bases of non-uniformities in spatial vision.

The area in visual space over which flanker stimuli impact the perception of a central target is referred to as the crowding zone. A fundamental property of crowding zones is their non-uniformity in visual space (for review, see *17*). Psychophysical studies have identified two major non-uniform characteristics. First, flanker stimuli positioned on the radial axis exert a larger influence on perception than flankers on the tangential axis (*18, Figure 1A*). Second, crowding is asymmetrical along the radial axis, such that flankers that are more eccentric than the target more strongly impair perception than flankers that are less eccentric than the target (*19*). Together, these results indicate that flanker stimuli positioned on the radial axis and more eccentric than the target (radial-out) are more effective at impairing target perception than stimuli either on the radial axis but less eccentric than the target (radial-in) or on the tangential axis (tangential).

**Figure 1.**
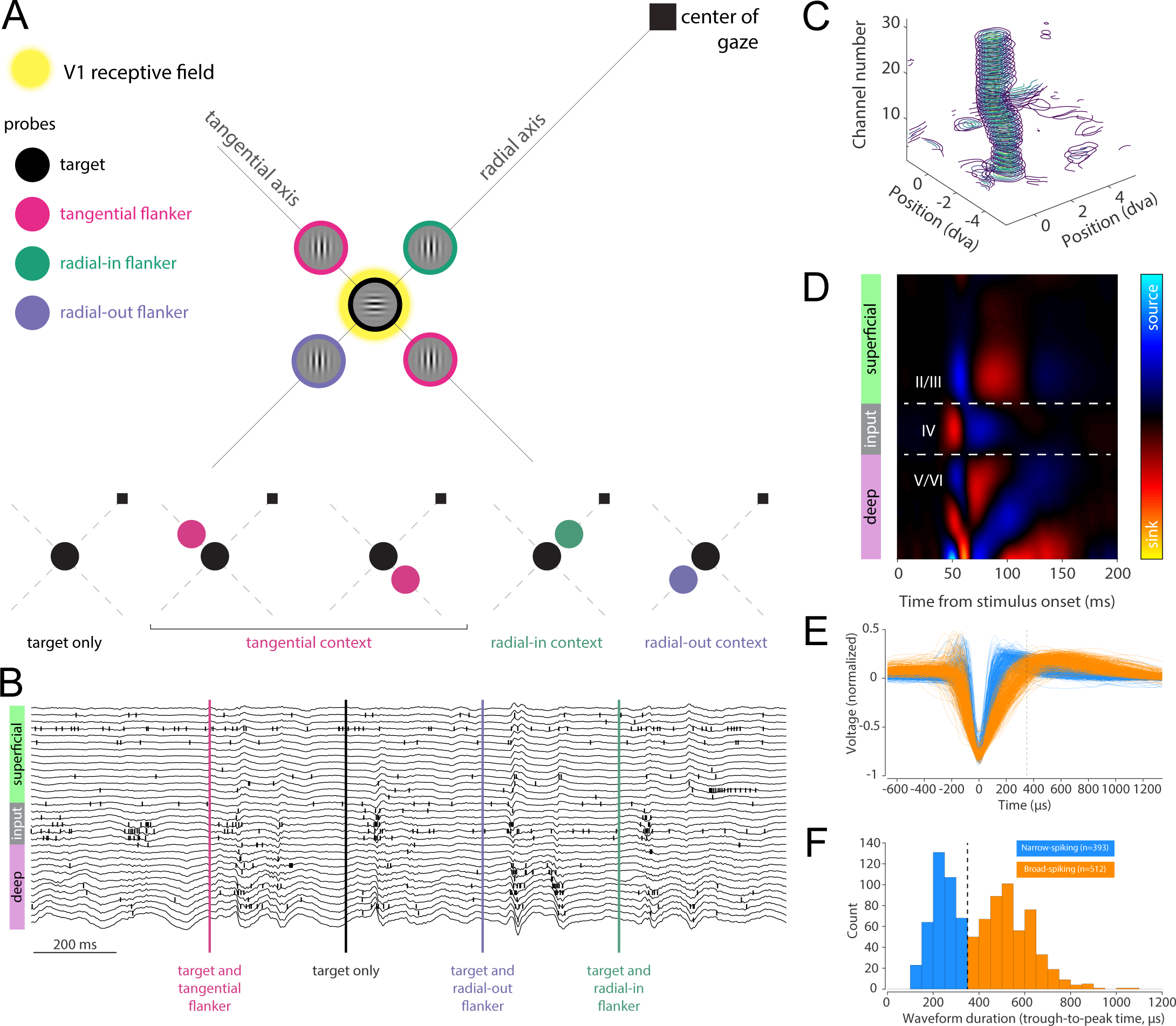
Behavioral task and single-unit recordings. (**A**) Task schematic. Stimuli were presented alone at the receptive field center (target only) or with a flanker at one of the four locations surrounding the target. The radial axis connects the target to the center of gaze, and the tangential axis is orthogonal to it, also passing through the target. The top schematic shows stimuli at all possible locations, while the bottom section shows the possible spatial configurations for the presentation of an individual stimulus array. (**B**) Data recorded from a single trial, including spikes (black vertical ticks) and local field potentials (black traces) collected from each channel on one of the two shanks of the laminar probe. Data is arranged vertically based on the spatial location of the recording channel. Cortical layers are indicated on the left (green=superficial, gray=input, pink=deep). Vertical bars indicate the time of stimulus onsets. (**C**) Example receptive field contours vertically stacked for each recording site along the depth of a shank on the laminar probe. (**D**) Example current source density (CSD) map showing the early current sink (red) indicative of the input layer. (**E**) Mean waveforms for all recorded single-units (blue=narrow-spiking, orange=broad-spiking). (**F**) Distribution of waveform durations (trough-to-peak time) for all recorded single-units across the two monkeys. Neurons with waveform durations less than 350µs were classified as narrow-spiking (blue), and neurons with waveform durations greater than 350µs were classified as broad-spiking (orange).

While the perceptual effects of crowding have been studied for many decades (*16, 17, 20–23*), it was only recently that there has emerged direct evidence of neural correlates underlying the phenomenon (*24, 25*). The question of exactly when and how crowding effects emerge in the visual hierarchy remains unresolved, but recent work has shown that crowding impairs feature representations as early as V1 (*24, 25*). However, the neural mechanisms underlying the spatial non-uniformities in perception remain poorly understood at the level of cortical microcircuits composed of multiple laminae and cell classes (*26–29*).

We sought to understand how stimuli presented outside the receptive field center could influence stimulus representations in the V1 laminar microcircuit in a spatially non-uniform manner. We hypothesized that contextual interactions with radial-out flankers would most strongly impact the representation of target stimuli among excitatory neurons in the output layers of V1. We further hypothesized that these non-uniform contextual interactions might be most prevalent in the subpopulations involved in feed-forward information propagation, particularly excitatory neurons in the input and superficial layers. These neurons play a key role in the propagation of stimulus information to higher cortical areas and are therefore critical for perception. Evidence of spatially non-uniform impairment of stimulus representations in these neural pathways would provide key insights into the circuit mechanisms in early visual processing that limit spatial vision.

## RESULTS

To study how the spatial configuration of stimuli influences neural responses in area V1, we trained monkeys to perform a passive fixation task while we presented stimuli in different geometric arrangements relative to each neuron’s receptive field. Monkeys fixated on the center of the screen and passively viewed stimuli presented either alone (target-only condition) or along with a flanker in one of four locations (context condition). We defined three context conditions based on the spatial position of the flanker relative to the target (Figure 1A, see Methods). The radial-in and radial-out context conditions positioned the flanker on the radial axis connecting the target to the fixation point. In the radial-in condition, the flanker was placed between the target and the fixation point, while in the radial-out condition, the flanker was past the target on the radial axis. For the tangential context condition, the flanker was positioned on the tangential axis (the line orthogonal to the radial axis and passing through the target) on either side of the target.

During the task, we performed laminar extracellular recordings (*30, 31*) of single unit activity in V1 through an artificial dura (Figure 1B-F, S1A-B). Probes were inserted orthogonal to the cortical surface to record from each layer along the depth of a column (Figure 1C-D). The receptive fields of all recording sites were well-aligned in visual space (Figure 1C). We positioned the stimuli such that the target stimulus was presented at the center of the aggregate receptive field of the recording sites. During our recordings, we collected data from 905 well-isolated single units (Figure 1E, 511 from monkey M, 394 from monkey D). We used established procedures (*30, 32–36*) to classify these neurons based on their waveform shape into broad-spiking (putative excitatory) and narrow-spiking (putative inhibitory) neurons (Figure 1E-F). Many observed differences between broad- and narrow-spiking neurons suggest they correspond well with excitatory and inhibitory neurons respectively (*30, 33, 34, 36–40*). We used current source density (CSD) analysis to identify the superficial (II/III), input (IV), and deep (V & VI) cortical layers (*30, 41*) (Figure 1D). This allowed us to analyze the different impacts of flanker stimuli based on their spatial location in six subpopulations of the laminar cortical circuit: broad- and narrow-spiking neurons in each of the three layers. Each of these subpopulations was visually responsive and well-tuned to the orientation of the target stimulus (Figure S2A-B & S4).

To examine how contextual interactions among stimuli might impact information coding in V1 in a non-uniform manner, we first determined how well we could decode the identity (orientation and phase) of the target, alone or in each of the context conditions, from simultaneously recorded V1 population activity using linear discriminant analysis (Figure 2). All flanker stimuli impaired decoding performance, with radial-out showing the largest performance decrease (Figure 2).

**Figure 2.**
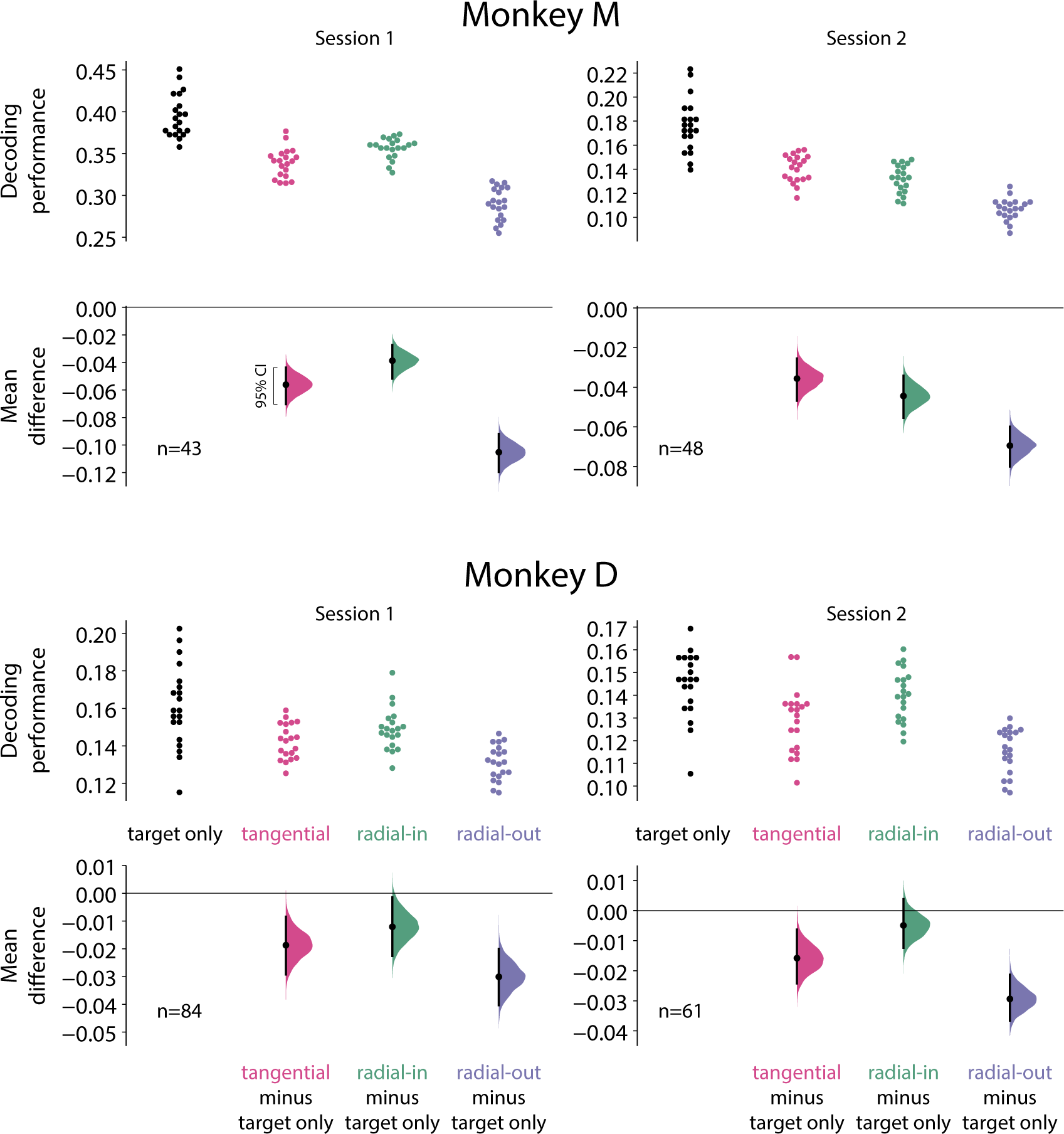
Flanker stimuli impair decoding performance in simultaneously recorded populations in a spatially non-uniform manner. Decoding performance from two example sessions in each monkey in the target-only condition and each of the three context conditions. Points in the upper section of each plot show the decoding performance for each of the 20 different cross-validations. The lower section for each session shows the bootstrapped estimation of the difference between decoding performance in each context condition compared to the target-only condition. Half-violin plots show the bootstrapped distribution of the difference, and black dots and bars represent the mean and 95% confidence intervals of the difference in decoding performance.

We next sought to understand how flanker location differentially influences information processing among laminar cortical circuit components. For our main analyses, we chose to focus on the broad-spiking populations, because these represent the primary output neurons of the column. For each subpopulation, we calculated the decoding performance for the target-only and context conditions (Figure 3; see Figure S3A for narrow-spiking populations). Most flanker stimuli impaired decoding performance compared to the target-only in most subpopulations (Figure 3 & S3A). In particular, the impairment of decoding performance was worse in the radial-out condition compared to the tangential and radial-in conditions among broad-spiking neurons in the input and superficial layers, the two subpopulations that are key nodes in the feed-forward cortical information flow pathway (Figure 3, middle and top). Surprisingly, tangential flankers more strongly impaired decoding in the deep layers (Figure 3, bottom). Radial-out flankers also impaired decoding performance more than radial-in flankers in narrow-spiking neurons across all layers, although radial-out flankers had a similar impact compared to tangential flankers in these subpopulations (Figure S3A). Consistent with previous reports (*25*), we also found that stimuli associated with stronger crowding effects (radial-out) most strongly reduced spike-count correlations, and this result was strongest in the feed-forward pathway (Figure S3B-C).

**Figure 3.**
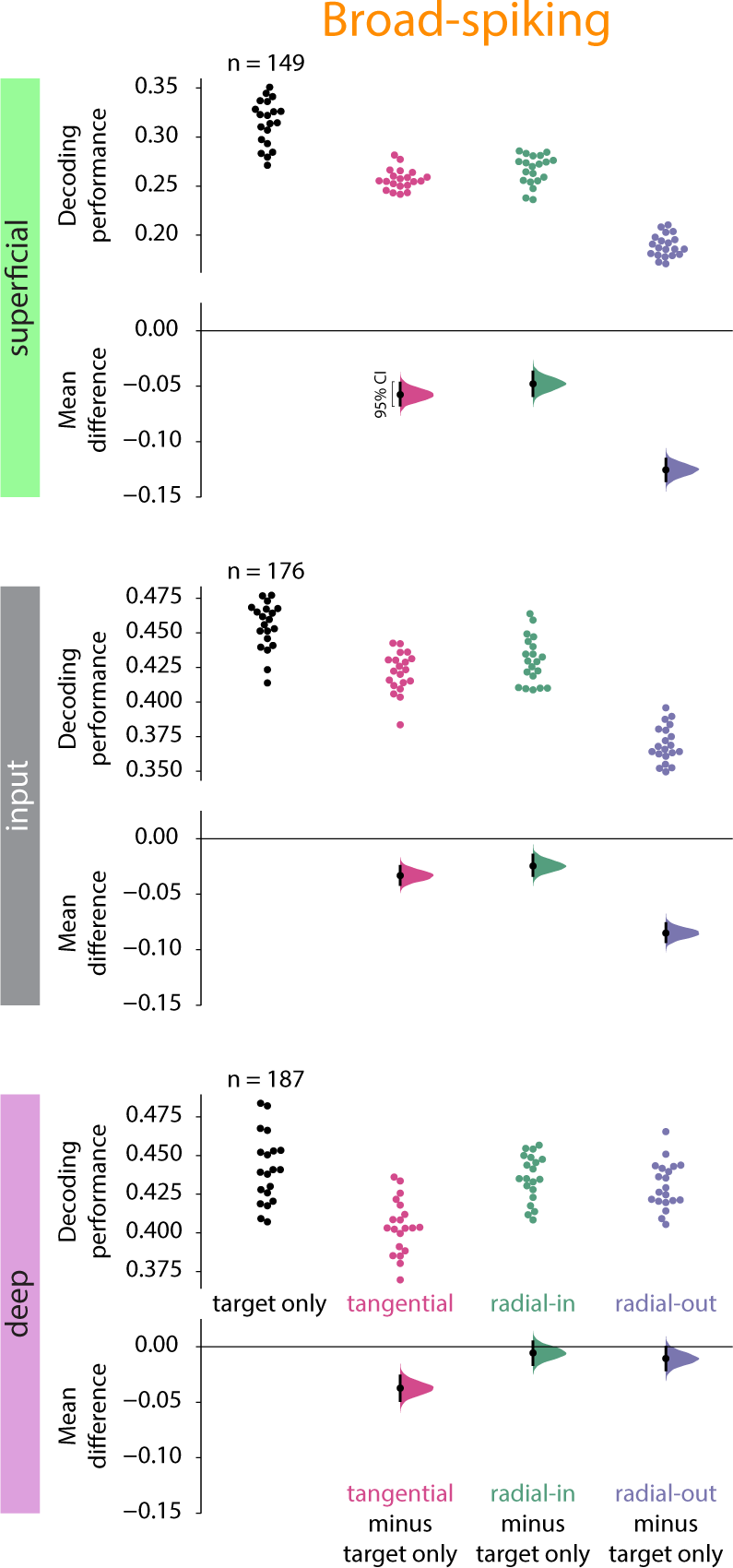
Radial-out interactions most strongly impair decoding in the feed-forward pathway. Decoding performance from broad-spiking neurons in each layer in the target-only condition and each of the three context conditions. Points in the upper section of each plot show the decoding performance for each of the 20 different cross-validations. The lower section for each layer shows the bootstrapped estimation of the difference between decoding performance in each context condition compared to the target-only condition. Half-violin plots show the bootstrapped distribution of the difference, and black dots and bars represent the mean and 95% confidence intervals of the difference in decoding performance.

Because flanker location has a non-uniform impact on the ability to decode target orientation from population activity, we investigated whether differential modulation in the tuning properties of V1 neurons could underlie these changes. We normalized the tuning curve for each single unit based on its preferred target orientation (see Methods) and plotted the average tuning curve of each subpopulation in response to the target only and in the context conditions (Figure 4A & S4A). Flankers broadened the tuning curves of V1 neurons in each of the subpopulations (Figure 4 & S4A). All subpopulations showed substantially reduced responses when flankers were presented alone (Figure S4C).

**Figure 4.**
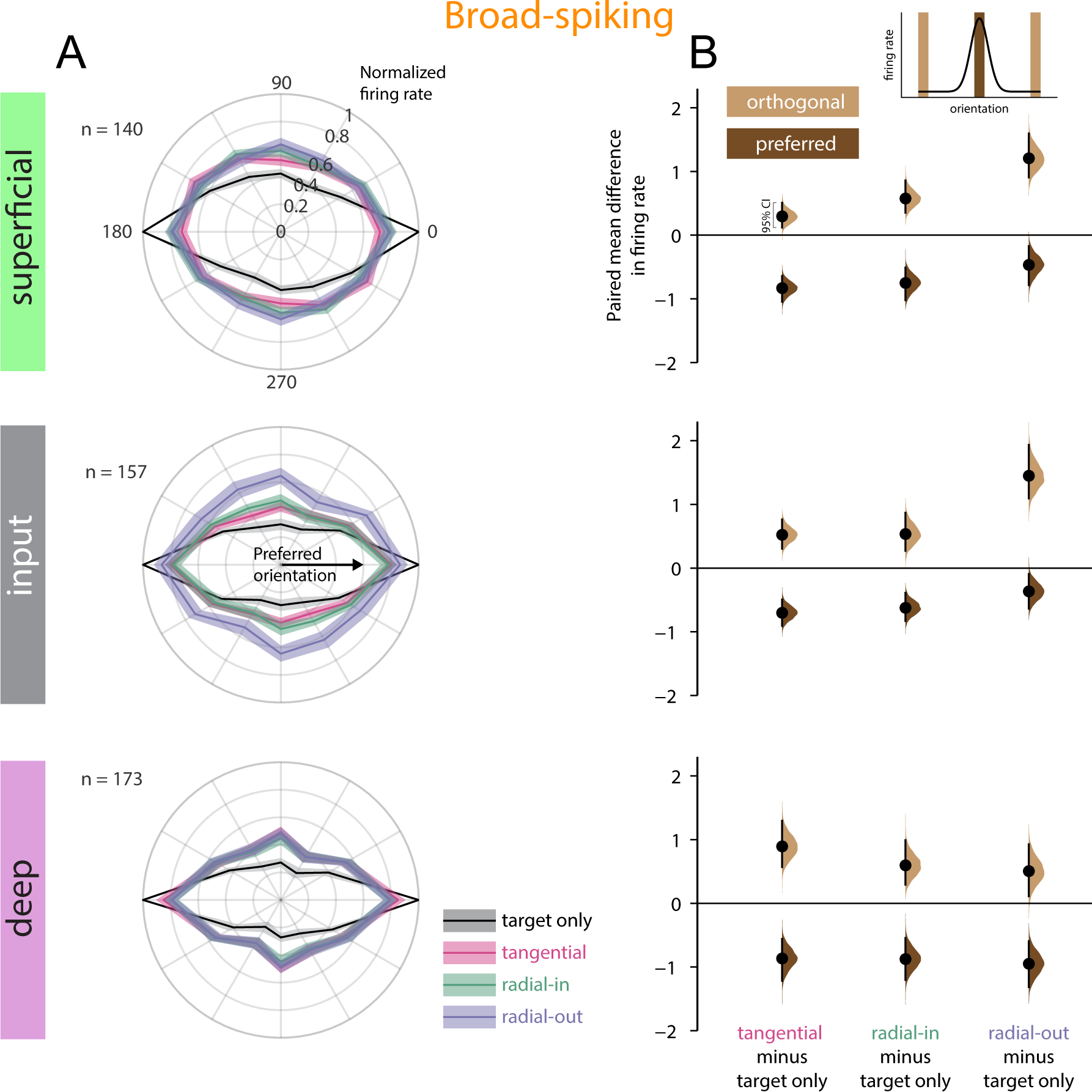
Flanker stimuli modulate the tuning curves of V1 neurons. (**A**) Normalized tuning curves for broad-spiking neurons in each layer in the target-only and context conditions. Tuning curves are aligned with the preferred orientation at 0 degrees and duplicated to fill the circle. Tuning curves were normalized by the maximum firing rate in the target-only condition. Lines and shading represent mean +/− s.e.m. (black=target only, pink=tangential context, green=radial-in context, purple=radial-out context). (**B**) Difference in firing rate in response to preferred (dark brown) and orthogonal (light tan) stimuli under each context condition across layers. Colored half-violin plots show the bootstrapped estimation of the paired mean difference in firing rate for a given subpopulation, context condition, and stimulus orientation. Black dots and lines represent the mean and 95% confidence interval of the estimated difference.

To quantify how flanker location modulated tuning, we first calculated the firing rates in response to preferred and orthogonal-to-preferred (orthogonal) target orientations. Flanker stimuli reduced firing rates in response to the preferred target orientation and increased the firing rates in response to the orthogonal target orientation (Figure 4B & S4B). This correspondingly resulted in a decrease in orientation selectivity index (OSI) that was similar across context conditions (Figure S2C). Although OSI was similar across conditions and subpopulations, there were distinct differences in how flanker location modulated preferred and orthogonal firing rates. In broad-spiking neurons in both the input and superficial layers, radial-out flankers resulted in less suppression of firing to the preferred orientation, but more effectively facilitated firing in response to the orthogonal orientation (Figure 4B, middle and top). This pattern was also consistent among narrow-spiking neurons in the input layer (Figure S4B, top). These differences were not due to changes in average eye position across conditions (Figure S4D).

To further characterize these results and obtain mechanistic insights into these changes in tuning properties, we quantified the extent to which flanker location induced untuned additive facilitation or tuned divisive suppression (*24, 25*) of V1 tuning curves. We modeled each neuron’s tuning curve in the target-only condition and then calculated the extent to which the responses to stimuli under each of the context conditions were best explained by additive and multiplicative shifts to the target-only tuning curve (Figure 5 & S5). Input and superficial layer broad-spiking neurons in the presence of radial-out flanker stimuli showed greater additive facilitation compared to tangential or radial-in conditions. (Figure 5A, middle and top). The radial-out condition also showed greater divisive suppression in broad-spiking neurons in both the input and superficial layers compared to the tangential and radial-in conditions (Figure 5B, top and middle). This effect was similar in narrow-spiking neurons in the input and superficial layers, with the radial-out condition showing more divisive suppression in both subpopulations compared to radial-in condition and trending towards more suppression compared to tangential condition (Figure S5B, top and middle).

**Figure 5.**
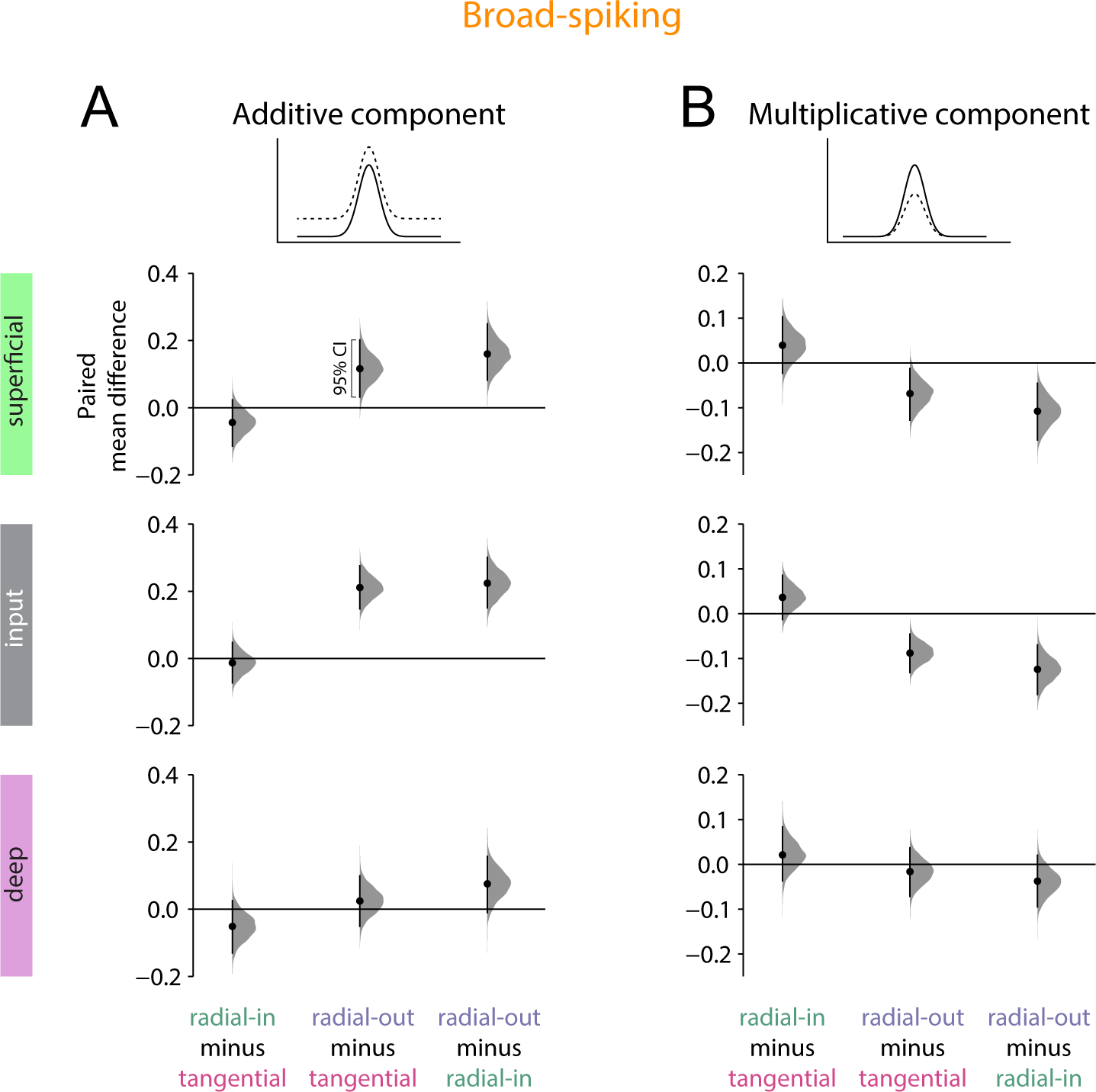
Radial-out contextual interactions cause stronger additive facilitation and divisive suppression of tuning curves. (**A**) Difference in the modeled (see Methods) untuned additive facilitation of tuning curves of broad-spiking neurons caused by each of the context conditions. Gray half-violin plots show the bootstrapped estimation of the paired mean difference in the modeled additive component for a given subpopulation and pair of context conditions. Black dots and lines represent the mean and 95% confidence interval of the estimated difference. (**B**) Same as (**A**) but for the multiplicative component, reflecting tuned divisive suppression.

We finally sought to understand how non-uniform integration could be explained in the context of normalization, a powerful explanatory framework for cortical computations (*42, 43*). We adapted the normalization model (*44*) by skewing the inhibitory kernel towards the fovea and introduced an additional additive kernel that was also skewed towards the fovea (Figure 6A, see Methods). For simplicity, visual space in the model is one-dimensional along the radial axis, and thus we chose to include only radial-in and radial-out context conditions. With these skewed kernels, orientation tuning curves for neurons with receptive fields at the target location were substantially broader in the radial-out condition than radial-in condition or when the target was presented alone (Figure 6B). The population response of neurons with receptive fields at the target location was more strongly disrupted by the radial-out flanker compared to the radial-in flanker (Figure 6C). Moreover, the tuning curves predicted by the model in the different context conditions qualitatively resembled the population-averaged tuning curves for broad-spiking neurons in the input layer (Figure 6D).

**Figure 6.**
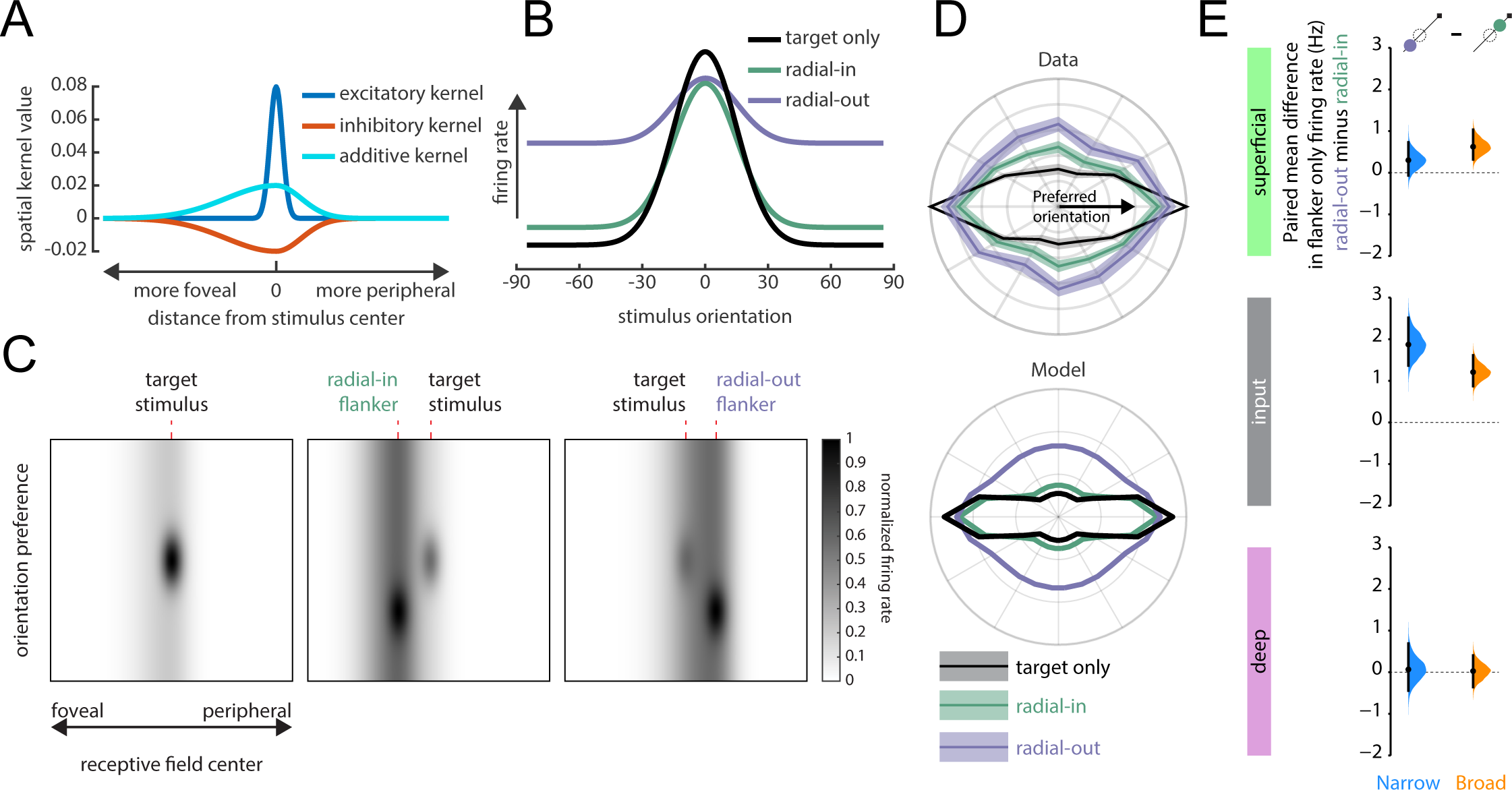
Normalization model with spatially skewed additive and divisive components explains non-uniform contextual interactions. (**A**) Excitatory (blue), inhibitory (red), and additive (cyan) kernels used in the normalization model. The additive and inhibitory kernels are skewed by doubling their standard deviation in the foveal direction and scaling both sides to maintain a smooth curve. (**B**) Modeled tuning curves in the target only (black), radial-in context (green), and radial-out context (purple). (**C**) Modeled population response in the three conditions: target only (left), radial-in context (middle), and radial-out context (right). (**D**) Comparison of population-averaged tuning curves from input-layer broad-spiking neurons (top, same as Figure 4A) and modeled tuning curves (bottom). (**E**) Difference in firing rate for each of the six subpopulations when a flanking stimulus was presented alone in the radial-out position compared to the radial-in position without the target present. Black dots and lines represent the mean and 95% confidence interval of the estimated difference.

The skewed normalization model suggests that stimuli should elicit stronger responses in neurons with receptive fields that are less eccentric compared to neurons with more eccentric receptive fields. This led us to predict that neurons with receptive fields at the target location should respond more to the radial-out flanker when presented alone (without the target stimulus) than to the radial-in flanker alone. Indeed, broad-spiking neurons in the input and superficial layers had higher firing rates when the flanker was presented at the radial-out location compared to the radial-in location (Figure 6E & S6), as did narrow-spiking neurons in the input layer (Figure 6E & S6). This analysis thus provides direct evidence of skewed kernels in visual cortex and confirms our model prediction that a stimulus would exert a stronger influence on neurons with less eccentric receptive fields even when presented alone.

Taken together, these results show that radial-out contextual interactions most effectively broadened tuning in broad-spiking neurons in the feed-forward pathway through untuned facilitation and tuned suppression. We propose this non-uniform interaction among stimuli could be explained by skewed additive and inhibitory interactions among neurons in V1.

## DISCUSSION

Psychophysical studies have suggested that crowding emerges in the cortex (*45, 46*), and various models (*21, 47–50*) have been proposed to explain crowding effects based on the anatomy and connectivity within the primary visual cortex (V1). Prior studies have hypothesized crowding can emerge due to properties in cortical areas V1 (*21, 51*), V2 (*52*), or V4 (*53*). Mechanistic explanations of crowding include averaging across stimuli (*54, 55*), inaccurately representing distractors as targets (*56*), or summary statistic representations of peripheral vision (*57*). Cortical magnification results in radial-out flankers being represented closer to the target in cortical space compared to radial-in flankers, potentially explaining some of the non-uniformity (*21, 48, 58*). However, it has been shown that considerations of cortical magnification alone are not sufficient to explain the marked elongation of the crowding zones along the radial axis defined by the center of gaze and the target stimulus (*47*).

While several studies (*59–61*) have explored the spatial distribution of contextual interactions and their impact on neural responses, systematic geometrical arrangements of target and flanker objects with respect to the center of gaze were not investigated. Furthermore, it has been shown that intrinsic horizontal connections in V1 are anisotropic in a manner that could facilitate contour processing, enriching connections among neurons with orientation preferences and receptive fields that facilitate the detection of smooth contours (*62, 63*). But here too the spatial geometries pertinent to prominent perceptual non-uniformities have not been investigated.

Macaques experience crowding similar to humans, impairing their ability to discriminate the orientation of target stimuli (*23, 24*), and are thus a good model for studying the neural mechanisms of the non-uniformities underlying spatial vision. Experiments in both macaques and humans suggest that crowding can influence neural activity as early as V1 (*24, 64*), including a radial-tangential anisotropy in BOLD responses in V1 (*65*). The present study extends these findings by revealing the impact of flanker location on stimulus representation in the laminar microcircuit of the primary visual cortex and the neural mechanisms underlying these impacted representations.

Notably, the effects of radial-out contextual interactions were more pronounced compared to the tangential and radial-in conditions across various measures. These effects included the ability to decode target identity from simultaneously recorded populations, as well as from putative excitatory neurons in the input and superficial layers. Non-uniformities were also evident in the modulation of tuning curves in these layers, with radial-out contextual interactions producing stronger broadening of tuning curves through untuned additive facilitation and tuned divisive suppression. Additionally, radial-out flankers exhibited the most substantial decrease in spike-count correlations in the input and superficial layers. While decreases in correlation are typically associated with increased information coding capacity, correlations are not necessarily information limiting, and decreased correlation can worsen the encoding capacity of a population (*66, 67*). These non-uniformities likely contribute to the impaired perception of target stimuli induced by radial-out flankers. Importantly, the decreased ability to decode stimulus orientation from neural activity at the subpopulation level was not solely attributable to changes in spike-count correlations, as correlations were removed across all conditions for those analyses.

Surprisingly, we discovered that contextual interactions exhibit non-uniformity even at the earliest stages of processing in V1, specifically within the input layer. This finding suggests a potential sub-cortical origin of the observed non-uniformities (*68–72*). Additionally, we observed the presence of non-uniformity in the superficial layer of V1, implying that these computations are partially inherited from the input layer. Broad-spiking neurons share electrophysiological characteristics with excitatory pyramidal neurons (*34, 36, 40*). Given that broad-spiking putative excitatory neurons in the superficial layer are likely projection neurons to downstream ventral visual areas (*26, 73–75*), our findings suggest that the non-uniform signaling is propagated along the feedforward pathway and could form the basis of the perceptual non-uniformities underlying visual crowding.

In contrast, we did not find the same non-uniformity in the deep layers of V1. This intriguing result suggests the emergence of novel contextual computations in the deep layers, indicating that tuning and response properties are not merely inherited from input and superficial layer projections to the deep layers. This is in agreement with previous findings that neurons in the deep layers can have differences in orientation tuning compared to the superficial layers (*76, 77*) and that intrinsic (horizontal) connections in the deep layers are less extensive compared to the superficial layers (*78*). Moreover, as deep-layer broad-spiking putative excitatory neurons are the likely sources of feedback projections to subcortical areas, these feedback signals are likely spatially uniform in nature. Despite the likely feedforward propagation of non-uniform signals, the absence of the same in the feedback pathway suggests a mechanism of active cancellation in this important modulatory pathway.

The implications of our findings extend to the integration of information from outside the classical receptive field in the visual cortex. Orientation tuning properties and their modulation are determined by the balance of excitation and inhibition to cortical neurons (*79, 80*). Recurrent excitation (*81*) and lateral inhibition (*82*) are two key intra-cortical motifs that determine this balance. Cross-orientation suppression (*83*), a form of lateral inhibition likely plays an important role in shaping the cortical activity due to flanker-target interactions examined in this study. Such suppressive activity may operate in several ways according to orientation dependence and the strength of inhibitory connections and have been described in various models of cortical activity including those with attractor dynamics (*84, 85*), recurrence (*81*), balance (*86–88*), push-pull (*89, 90*) and normalization (*91–93*). Our results imply that flanker location differentially impacts the balance of excitation and inhibition in the laminar cortical circuit. Normalization, in particular, has been proposed as a canonical computation in the cortex (*43*) that explains a wide range of cortical phenomena including contrast response properties of visual neurons (*92, 94, 95*), multi-sensory integration (*96*), and multiple forms of attentional modulation (*44*) while remaining agnostic to the exact nature of the underlying biophysical mechanisms referred to above. Our modeling and empirical results show that the fundamental assumption of symmetry in the spatial kernels of the normalization model must be broken to account for the non-uniform nature of contextual integration in the visual cortex. Future empirical and computational modeling studies of surround modulation should take this non-uniform and layer-specific integration into account.

In conclusion, this study provides compelling evidence that contextual interactions in the visual cortex exhibit non-uniformity at the earliest stages of cortical processing of visual signals. Flankers broaden V1 orientation tuning curves in a spatially non-uniform manner. Radial-out flankers most effectively broaden tuning in the feedforward pathway through untuned facilitation and tuned suppression. These findings advance our understanding of the neural bases of spatial non-uniformities in perception and offer mechanistic insights into the processing of contextual signals in the visual cortex.

## AUTHOR CONTRIBUTIONS

MPM and ASN conceptualized the project. MPM collected the data, with assistance from SD and NVH. MPM analyzed the data. ASN supervised the project. MPM and ASN wrote the paper.

## ACKNOWLEDGEMENTS

We would like to thank Steve Chang, Weikang Shi and Alec Sheffield for their helpful comments on the manuscript. This research was supported by NIH/NEI R01 EY032555, NARSAD Young Investigator Grant, Ziegler Foundation Grant, and Yale Orthwein Scholar Funds to ASN, NIH/NINDS training grants T32-NS007224 and T32-NS041228 to MPM, and by an NIH/NEI core grant for vision research P30 EY026878 to Yale University. We would like to thank the veterinary and husbandry staff at Yale for excellent animal care.

---

**Figure S1.**
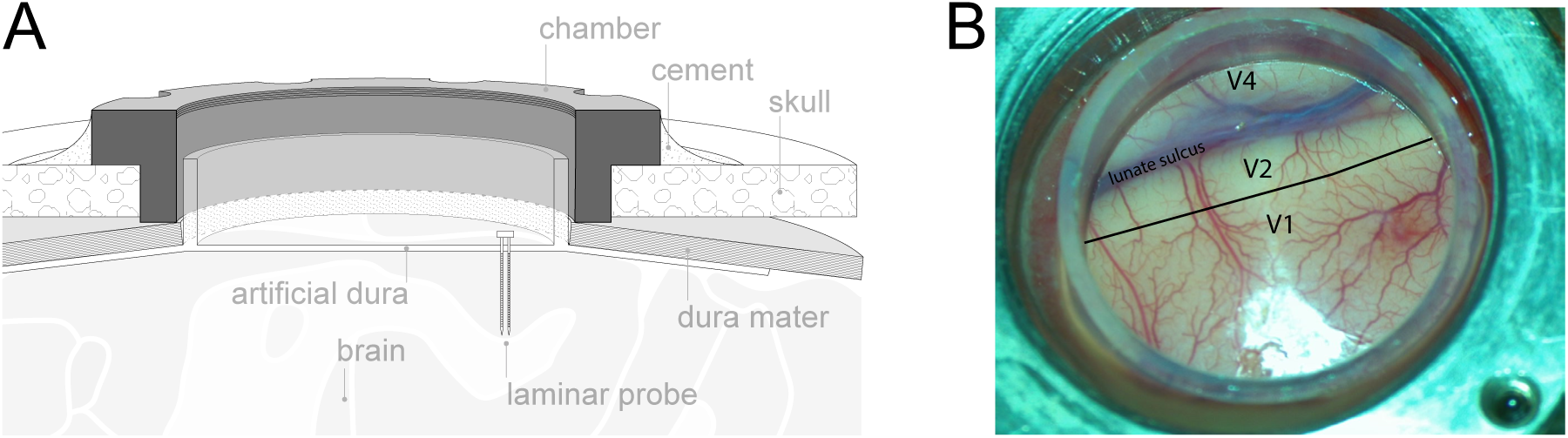
Laminar recordings in V1. (**A**) Section perspective schematic of the artificial dura recording chamber and laminar probe. (**B**) Photo of the artificial dura chamber in one monkey.

**Figure S2.**
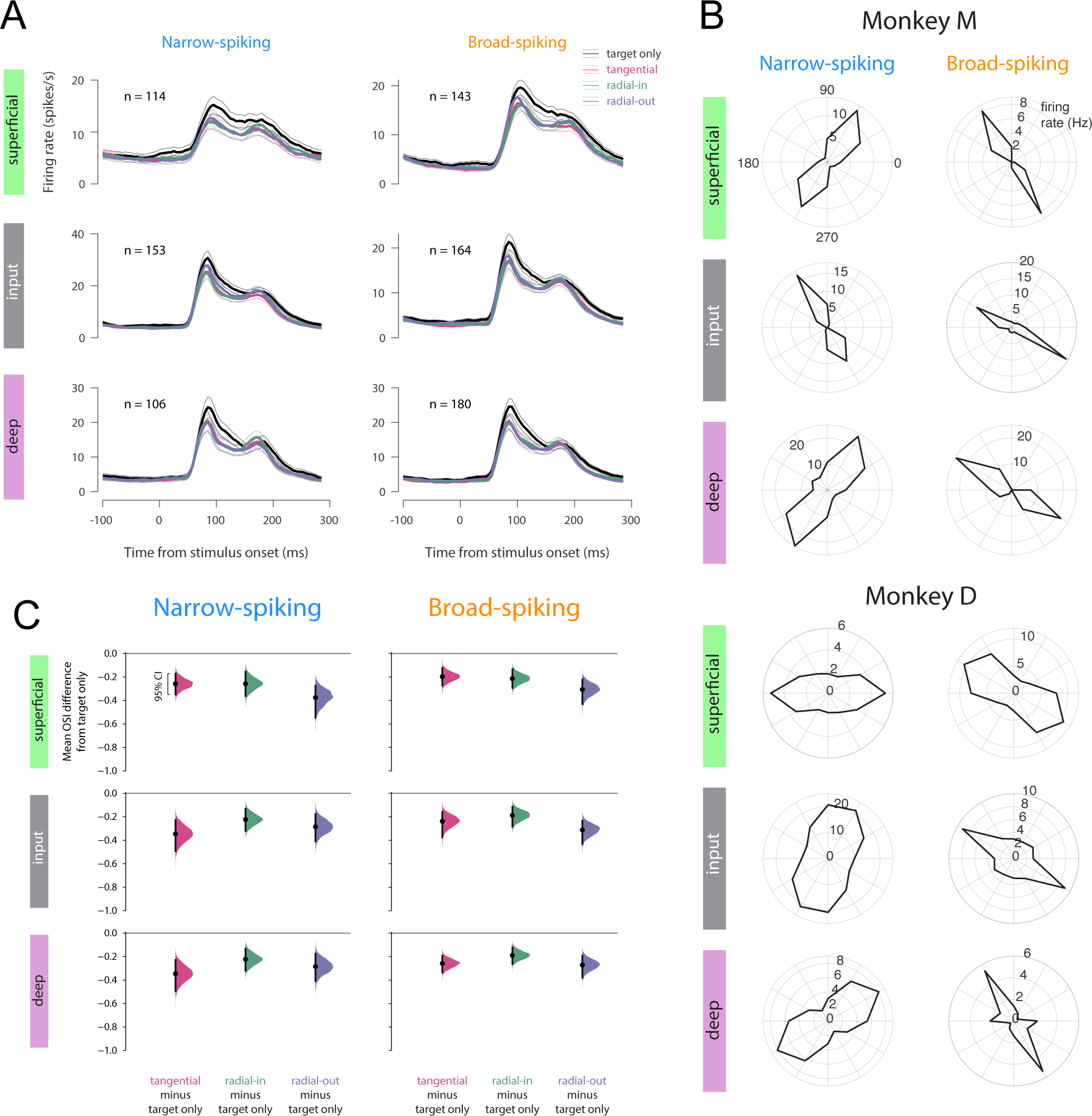
Peri-stimulus time histograms, tuning curves of single units, and orientation selectivity index under contextual interaction. (**A**) Peri-stimulus time histogram (PSTH) of subpopulations in each condition following the onset of the preferred stimulus. The preferred orientation was calculated separately for each single unit. Spikes were counted in 30ms bins shifted by 5ms. Error bars show mean+/− s.e.m. (**B**) Example single unit tuning curves in the target-only condition. Single units were selected from multiple different recording sessions. (**C**) Difference in orientation selectivity index (OSI) in each of the context conditions compared to the target-only condition. Colored half-violin plots show the bootstrapped estimation of the mean difference in OSI for a given subpopulation and context condition. Black dots and lines represent the mean and 95% confidence interval of the estimated difference.

**Figure S3.**
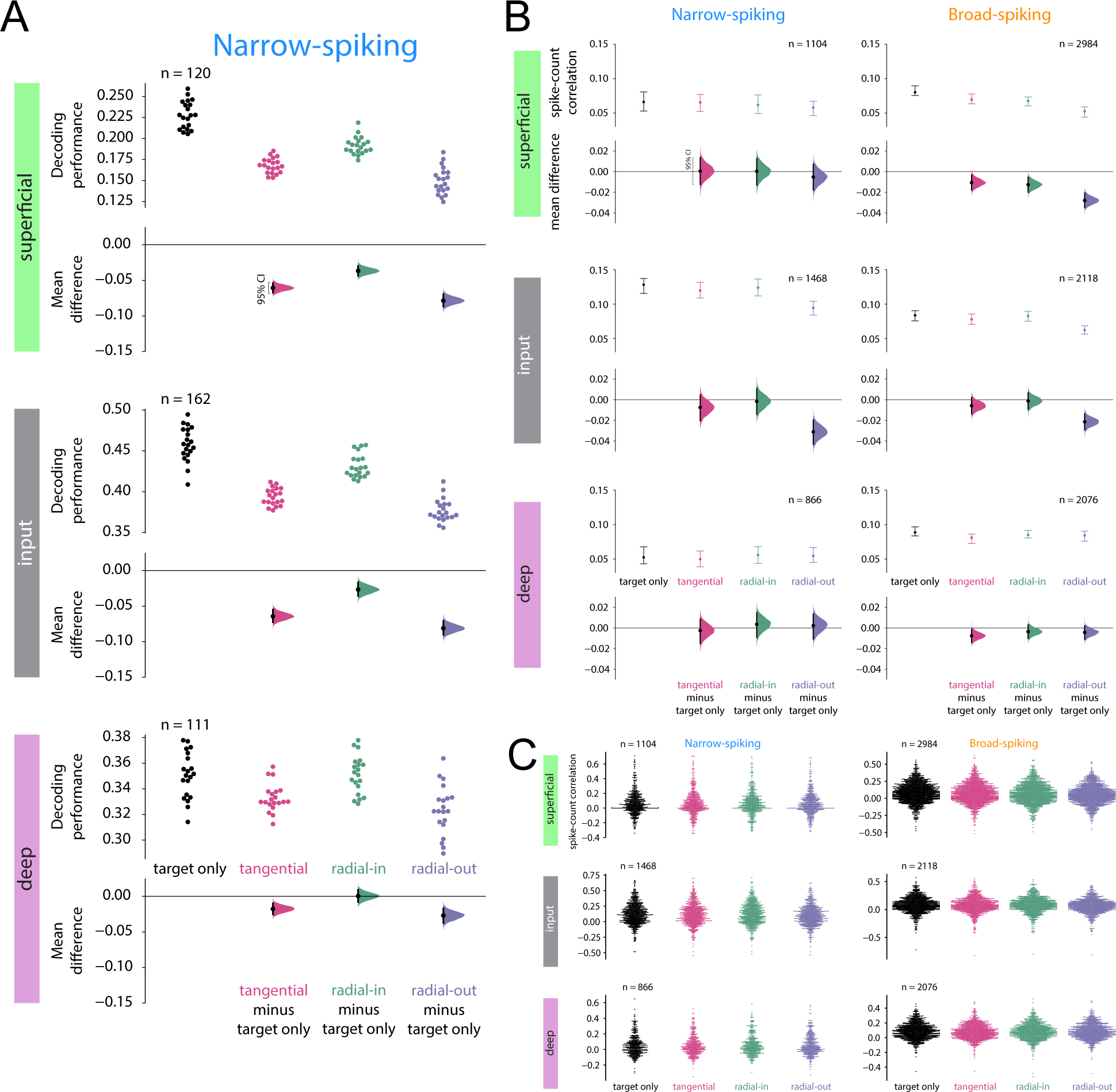
Decoding performance in narrow-spiking neurons and spike-count correlations. (**A**) Decoding performance from narrow-spiking neurons in each layer in the target-only condition and each of the three context conditions. Points in the upper section of each plot show the decoding performance for each of the 20 different cross-validations. The lower section for each layer shows the bootstrapped estimation of the difference between decoding performance in each context condition compared to the target-only condition. Half-violin plots show the bootstrapped distribution of the difference, and black dots and bars represent the mean and 95% confidence intervals of the difference in decoding performance. (**B**) Spike-count correlations from pairs of simultaneously recorded neurons in each subpopulation in the target-only condition and each of the three context conditions. In the top panels, error bars represent mean +/− 95% confidence intervals of the spike-count correlation in each condition. In the bottom panels, black dots and lines represent the mean and 95% confidence interval of the estimated difference from the target-only condition, and colored half-violin plots show the bootstrapped estimation of the mean difference in spike-count correlation for a given subpopulation and context condition. (**C**) Spike-count correlations for each pair of single units in each subpopulation. Each point represents the spike-count correlations of one pair of simultaneously recorded neurons in the same layer and cell class and each condition.

**Figure S4.**
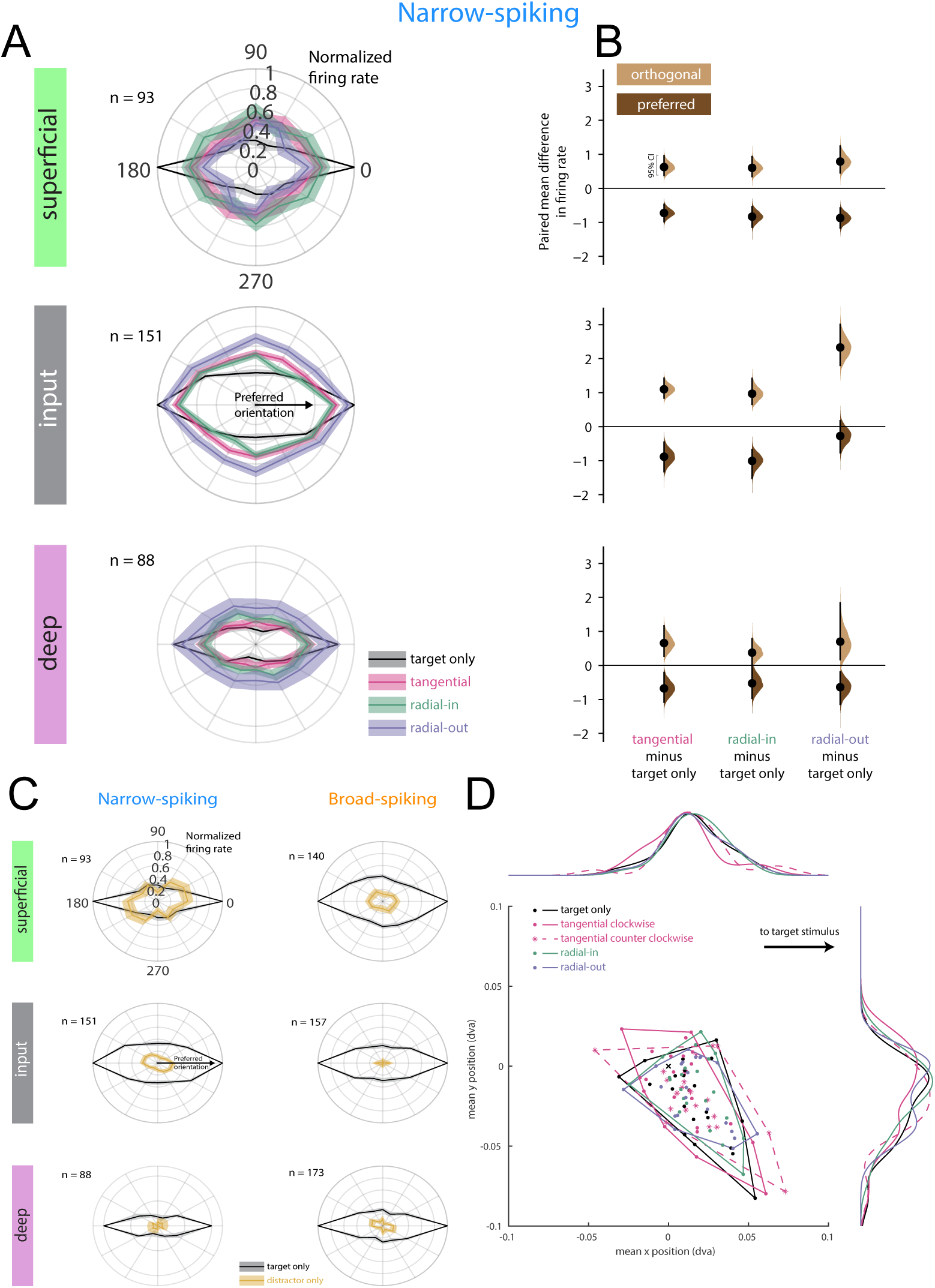
Tuning curves of narrow-spiking neurons, responses to flanker only, and eye position data. (**A**) Normalized tuning curves for narrow-spiking neurons in each layer in the target-only and context conditions. Tuning curves are aligned with the preferred orientation at 0 degrees and duplicated to fill the circle. Tuning curves were normalized by the maximum firing rate in the target-only condition. Lines and shading represent mean +/− s.e.m. (black=target only, pink=tangential context, green=radial-in context, purple=radial-out context). (**B**) Difference in firing rate in response to preferred (dark brown) and orthogonal (light tan) stimuli under each context condition among narrow spiking neurons in each layer. Colored half-violin plots show the bootstrapped estimation of the paired mean difference in firing rate for a given subpopulation, context condition, and stimulus orientation. Black dots and lines represent the mean and 95% confidence interval of the estimated difference. (**C**) Normalized tuning curves for each subpopulation in the target-only and flanker-only conditions. Tuning curves are aligned with the preferred orientation at 0 degrees and duplicated to fill the circle. Tuning curves were normalized by the maximum firing rate in the target-only condition. The flanker-only condition is plotted relative to the orientation of the flanker and combines data from all four flanker positions. Lines and shading represent mean +/− s.e.m. (black=target only, gold=flanker only). (**D**) Mean eye position during the four conditions in each recording session. Each session has five points plotted (target only, radial-in context, radial-out context, and tangential clockwise context as dots, tangential counter-clockwise as stars). Above and to the right are marginal distributions of the median x and y positions for each of the sessions in each condition (tangential counter-clockwise plotted as a dashed line). To account for different target positioning across sessions, all data are rotated such that the target stimulus is to the right.

**Figure S5.**
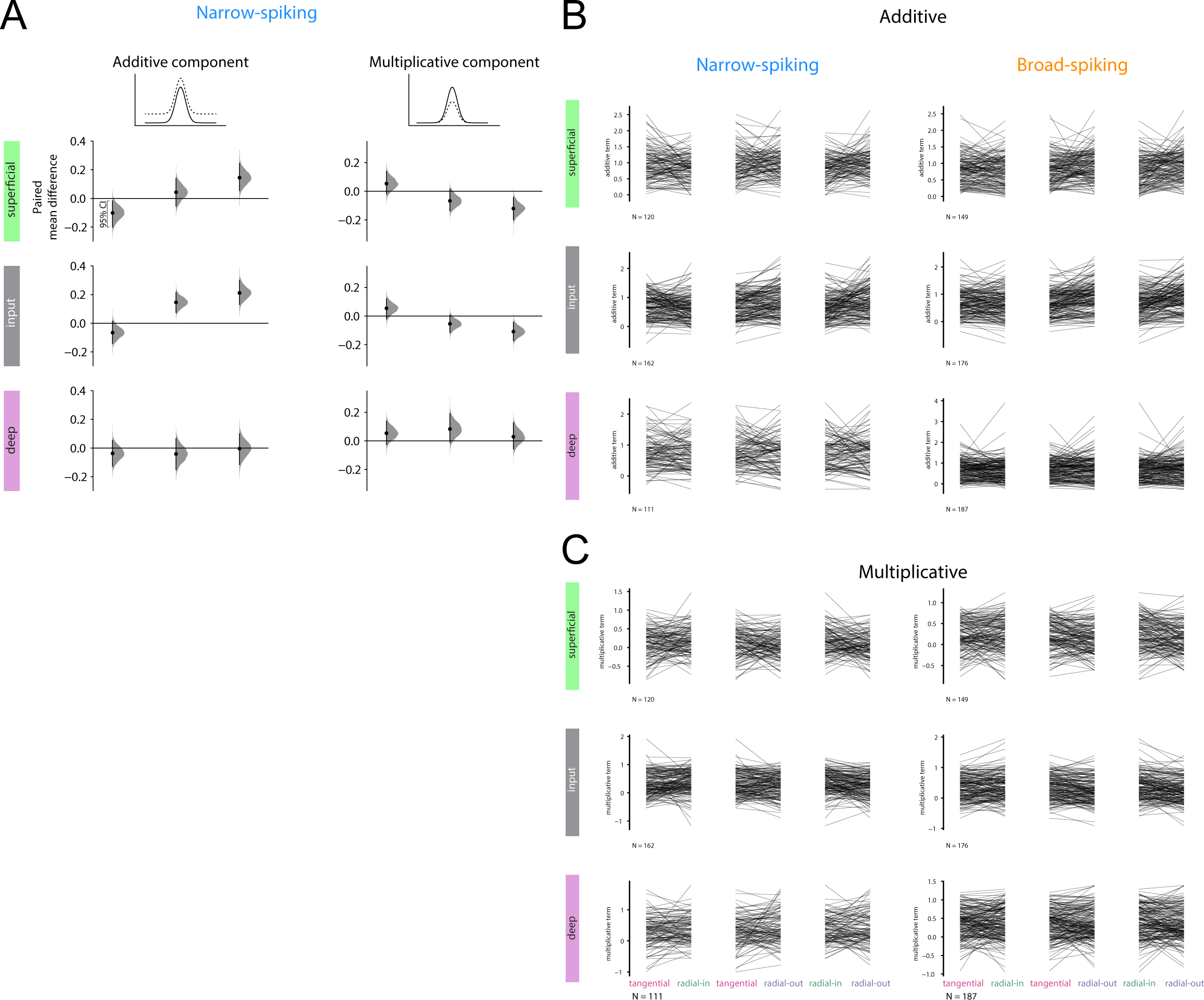
Single-unit tuning model parameters in different context conditions. (**A**) Difference in the modeled (see Methods) untuned additive facilitation (left) and multiplicative suppression (right) caused by each of the context conditions among narrow-spiking units. Gray half-violin plots show the bootstrapped estimation of the paired mean difference in the modeled additive component for a given subpopulation and pair of context conditions. Black dots and lines represent the mean and 95% confidence interval of the estimated difference. (**B**) Related to Figure 5A and S5A left. Value of the fitted additive parameter for each neuron in each context condition. Each line represents the additive parameter for a single unit in across the two context conditions indicated at the bottom. (**C**) Related to Figure 5B and S5A right. Same as (**B**) but for the multiplicative component.

**Figure S6.**
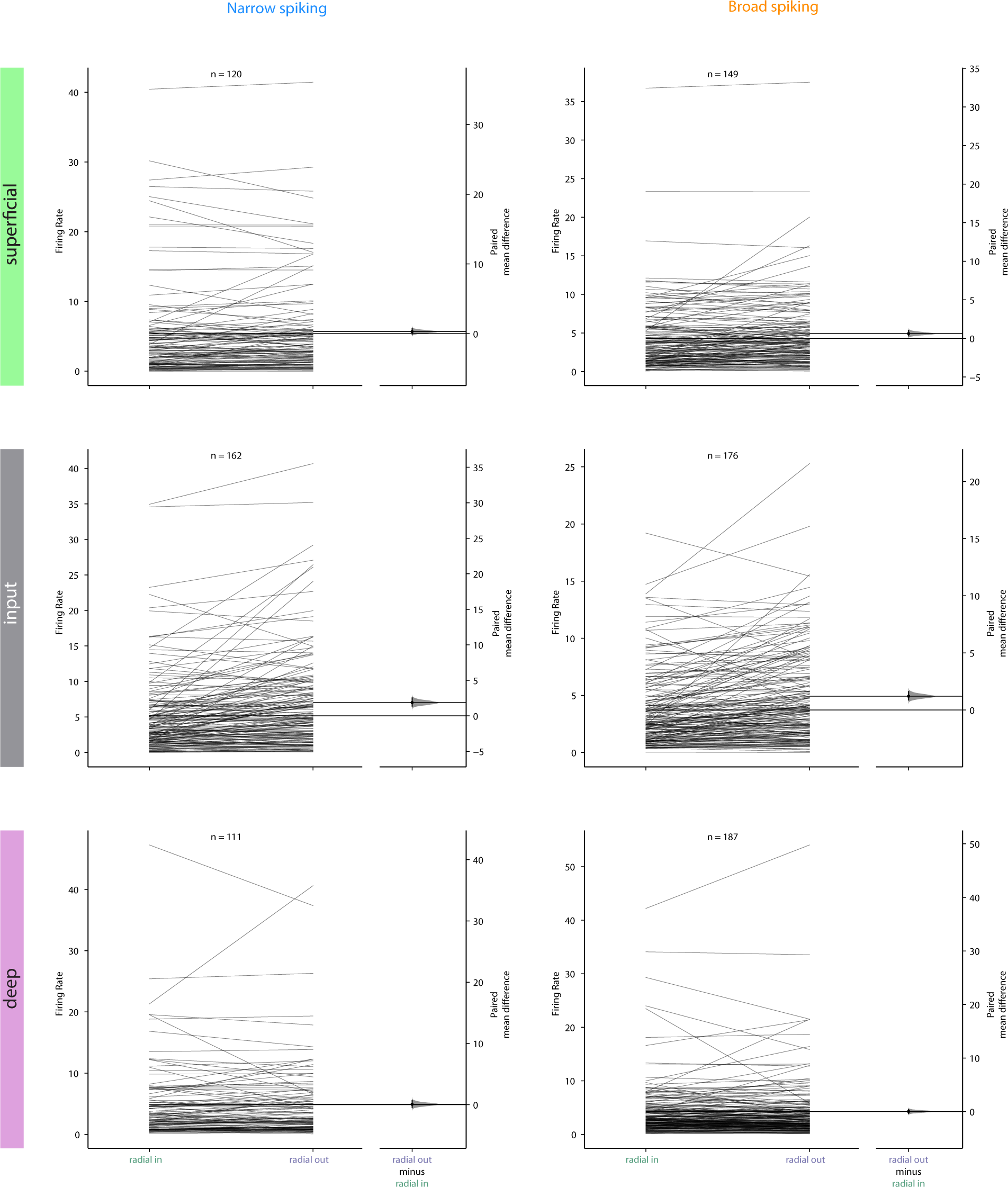
Single-unit firing rates in response to flanker alone. The average firing rate response of each neuron to stimuli in the radial-in or radial-out position is shown on the left side of the plot for each subpopulation. On the right, gray half-violin plots show the bootstrapped estimation of the paired mean difference in firing rate for a given subpopulation and pair of context conditions. Black dots and lines represent the mean and 95% confidence interval of the estimated difference. Estimation statistics plots are the same as those in Figure 6E but scaled based on the raw firing rates shown here so that axes on the left and right are identical.

## MATERIALS & METHODS

### Data Collection

#### Surgical procedures

Surgical procedures were similar to those described previously (*30, 31, 97*). We placed low-profile titanium recording chambers in two rhesus macaques so that the chambers allowed access to area V1 (right hemisphere in monkey D, left hemisphere in monkey M). Chambers were targeted based on sulcus reconstructions created using preoperative structural MRI. After chamber implantation, we removed the native dura mater and replaced it with a transparent silicone artificial dura (AD). The AD allowed for the visualization of cortical sites in V1 for probe targeting. All procedures were approved by the Yale University Institutional Animal Care and Use Committee and conformed to NIH guidelines.

#### Electrophysiology

Prior to a series of recordings, we electroplated (nanoZ, White Matter LLC) 64-channel electrode arrays (“laminar probes,” NeuroNexus Technologies, Inc., 2 shanks, 32 channels/shank, 70µm spacing between sites, 200µm between shanks) with PEDOT (poly(3,4-ethylene dioxythiophene)). At the beginning of each recording session, we inserted a laminar probe in V1. Laminar probes were attached to a titanium mounting stage that was screwed into the chamber. We positioned the laminar probes using an electronic micromanipulator (Narishige Inc.) and ensured that the probes were orthogonal to the surface of the cortex by visual inspection through a surgical microscope (Leica Microsystems). To position the probe within the brain, we first penetrated the AD, arachnoid, and pia by moving the probe downward at a high speed (>100µm/s). After the tip of the probe entered the cortex, we inserted the remainder of the probe at a slow speed (2µm/s). Once the entire probe was in the brain, we slowly (2µm/s) relieved the pressure on the brain by retracting the probe upward, relieving pressure on the brain but not moving the probe relative to the cortex.

Electrical signals from the laminar probe were collected at 30kHz and digitized on a 64-channel digital headstage and sent to the recording system (RHD Recording System, Intan Technologies). Action potential waveforms were extracted offline using Kilosort2 (*98, 99*) and manually sorted into single and multi-unit clusters (Phy; *98, 99*). Multi-unit clusters were excluded from further analysis. Single-unit clusters were classified based on their trough-to-peak waveform duration into broad- and narrow-spiking units (*30, 32*). Neurons with waveform durations less than 350µs were classified as narrow-spiking while those with waveform durations longer than 350µs were classified as broad-spiking. Clusters with peaks preceding the trough were identified as axonal spikes and excluded. Recordings were collected over the course of 22 sessions (9 in monkey D, 13 in monkey M) with hundreds of single units recorded in each subject (384 in monkey D, 521 in monkey M).

#### Behavioral Control and Eye Tracking

We controlled behavioral experiments using NIMH MonkeyLogic (*100*). Eye position and pupil diameter were sampled at 120Hz (ETL-200, ISCAN Inc.) and sent to the behavioral control system. Stimuli were presented on a monitor positioned 57cm from the monkey with a 60Hz refresh rate. Trials were aborted if the eye position deviated more than 1.2-1.5 degrees of visual angle (dva; 1.2 for monkey D, 1.5 for monkey M) from the central fixation point.

#### Receptive Field Mapping

We mapped RFs of the column by presenting Gabor patch stimuli (2-4 cycles/degree, 0.25-1 degree Gaussian half-width, 100% luminance contrast) on a square grid spanning the lower visual quadrant of interest (left in monkey D, right in monkey M) while the monkey maintained fixation on the center of the screen. Grid spacing parameters were optimized each session and ranged from 0.25-1 dva. A stimulus was presented at a random location and orientation on the grid during each frame. We calculated the LFP power for each recording channel 40-200ms after stimulus presentation in each location. The LFP power at each location was smoothed using a Gaussian kernel (sigma = 0.75 dva), and the peak location averaged across all recording sites was defined as the RF center. Spatial RF maps for each channel were plotted as stacked contours for each shank for visualization. The RF center was determined from the aggregate RF across all shanks and channels.

#### Current Source Density Mapping

We used CSD mapping (*41*) to identify laminar boundaries in our recordings. While monkeys maintained fixation on the screen, 100% luminance contrast white annular stimuli were flashed for 32ms, positioned at the center of the RF. The CSD was calculated as the second spatial derivative of the LFP following stimulus onset. CSD traces were spatially smoothed using a Gaussian kernel (sigma = 140µm). The input layer was identified by the boundaries of the early current sink characterizing feed-forward input into layer IV. Channels above and below this sink were classified as superficial and deep respectively.

#### Behavioral Task

While monkeys maintained fixation, stimulus arrays were flashed on the screen for 100ms and off the screen for 200ms. The target location was at the center of the aggregate RF of all channels. The arrays consisted of a target stimulus in the RF center either in isolation (target only) or together with a flanker stimulus (context). There were four different conditions (target only, tangential context, radial-in context, and radial-out context) of stimulus arrays presented during a trial. This was repeated 4-6 times during each trial depending on the monkey’s ability to hold fixation. The stimulus conditions were randomly interleaved from flash to flash. The center of the target stimulus was aligned with the center of the average of the RFs of each recording site. The four flanker locations were positioned on the radial or tangential axes. The radial axis was defined as the axis connecting the target (and RF) center and the fixation point at the center of the monitor. The tangential axis was defined as the line orthogonal to the radial axis and passing through the target center. The tangential context condition occurred when the target was presented along with a flanker stimulus on the tangential axis, either in the clockwise or counterclockwise direction. The radial-in context condition occurred when the target was presented along with a flanker stimulus on the radial axis and between the target center and the fixation point. The radial-out condition was similar to the radial-in condition, except the flanker stimulus was placed on the radial axis further from the fixation point than the target. Target-flanker spacing was identical across stimulus conditions.

The target-only condition was presented on 10% of flashes. On 80% of flashes, the context conditions were presented. In the context conditions, the flanker stimulus was positioned at a tangential, radial-in, or radial-out location with equal probability. If the flanker was presented tangential to the target, it was positioned clockwise or counterclockwise to the target with equal probability. In the remaining 10% of trials, a stimulus was presented exclusively at one of the four flanker locations. Target stimuli were sine Gabor patches (25% luminance contrast, 3.5 cycles/degree, 0.5-1.0 degree Gaussian half-width) presented at 6 evenly spaced orientations and 2 opposing phases. Flanking stimuli were presented at 100% luminance contrast but were otherwise identical to the target stimuli. There was a 0.1-0.2 degree gap between the edges of the target and flanking stimuli (edge defined as 2 standard deviations from the center of the Gaussian). In the context condition, flankers were presented at the same orientation as the target or orthogonal to the target with equal probability. In this study, we focused exclusively on the condition in which the flanker was orthogonal to the target. When flankers were presented alone, they were shown with the same orientation distribution as the targets.

### Data Analysis

Wherever possible, we analyzed our data using the estimation statistics framework (*101, 102*) and show bootstrapped estimations of differences between conditions rather than relying on null-hypothesis testing. The estimation statistics framework provides a principled way to measure effect sizes coupled with estimates of uncertainty, yielding interval estimates of uncertainty.

#### Tuning Curve Visualization

To visualize the modulation of tuning curves of each subpopulation by flanker stimuli, we calculated the number of spikes for each neuron 0-300ms following stimulus onset in each orientation and condition, collapsing across phases, and subtracted the baseline firing rate (0-100ms before stimulus onset). For each neuron, we identified the orientation with the highest firing rate in the target-only condition as the preferred orientation and shifted the tuning curves to set this orientation to 0 degrees in each condition. We then normalized the turning curves by dividing each neuron’s response to each orientation and condition by the firing rate for the preferred target-only orientation. For visualization, the tuning curve was duplicated and rotated 180 degrees to completely fit around a circle. We excluded neurons that did not respond in the target-only condition. To visualize the tuning curves of individual single units, we followed the same procedure but did not average across phases, instead plotting the response to each phase as the opposite orientation. Additionally, single-unit tuning curves were not rotated based on the preferred orientation.

#### Preferred and Orthogonal Firing Rate and OSI

OSI was calculated using the following equation:

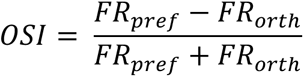

Where *FR_pref_* is the firing rate in response to the preferred orientation and *FR_orth_* is the firing rate in response to the stimulus orthogonal to the preferred orientation. Firing rates were calculated 0-300ms following stimulus onset. The preferred orientation was identified as the orientation with the highest firing rate. *FR_pref_*, *FR_orth_*, and OSI were compared between the target-only condition and the context conditions by calculating the bootstrapped estimation of the mean difference.

#### Modeling Tuning Curve Modulation

We investigated the components of single-unit tuning curve modulation using a previously published approach to identify additive and multiplicative changes to tuning curves (*25*). Briefly, we fit a Poisson model based on spike counts from 0-300ms after stimulus onset in the target-only condition for each orientation. We then calculated the untuned additive and tuned multiplicative modulation of the fitted target-only tuning curve that would best account for the observed spike counts independently for each context condition (*25*). We then compared the additive and multiplicative parameters for each neuron across conditions by calculating a bootstrapped estimation of the paired mean difference of the components.

#### Decoding Analysis

We trained a decoder using linear discriminant analysis (LDA) to determine the identity (orientation and phase) of a target stimulus from simultaneously recorded single-unit activity. Spike counts were calculated from 0-300ms following stimulus onset. The decoder performance was determined using 20-fold cross-validation. The decoder was trained on stimuli presented in the target-only condition and tested on neural responses in each of the context conditions. This procedure was repeated for each recording session. Performance in the tangential context condition was calculated separately for the two tangential locations and averaged.

To calculate decoding performance at the subpopulation level, we first generated a database of pseudo-population activity for each neuron in every combination of conditions and target orientations. Spike counts were calculated from 0-300ms following stimulus onset. To train the decoder for each subpopulation, we generated pseudo-population activity by randomly drawing samples from this database, independently for each neuron. We trained separate decoders for each subpopulation, training and testing on responses from every neuron in each cell class and layer across all recording sessions. Decoders were trained by sampling from data in the target-only condition. 20% of each neuron’s responses to each orientation were held out (the subset was independent for each neuron) for cross-validation. This procedure was repeated 20 times with a different random subset of responses used for cross-validation in the target-only condition. The decoding performance on this 20% of left-out data was considered the decoding performance in the target-only condition. We then tested each decoder on pseudo-population activity generated from responses in the context conditions.

#### Peri-stimulus time histograms (PSTH)

PSTHs were generated based on spike counts in 30ms bins shifted by 5ms. PSTHs are shown only for the preferred orientation as defined previously. The PSTH for each neuron was calculated separately for each condition, and the average PSTH for each subpopulation was plotted in Figure S2A. All times are relative to stimulus onset.

#### Spike-count correlations

Spike-count correlations were calculated for each pair of simultaneously recorded neurons in the same layer and broad- or narrow-spiking cell class. Spike counts were extracted from 100-300ms after stimulus onset and normalized by Z-score for each stimulus orientation and condition. For each condition, the spike-count correlation was calculated separately for each orientation from the normalized spike counts and was then averaged across orientations. Thus, each pair of neurons had a single spike-count correlation number for each stimulus condition.

### Eye-position analysis

In each session, we calculated the monkey’s average eye position in the time period from 50ms before to 150ms after each stimulus presentation. We then averaged these positions for each stimulus category: target only, radial-in context, radial-out context, and tangential (clockwise and counterclockwise) context within each session to obtain one value per condition and session. To compare across all sessions, we rotated the mean eye position values based on the location of the target stimulus so that the target was always to the right in Figure S4D.

### Normalization model

Our model was based on the normalization model presented by Reynolds and Heeger (*44, their Figure 1*). We excluded any attentional effects by making the attentional field uniform across both space and orientation. To make our kernels asymmetrical, we doubled the standard deviation of the Gaussian for the inhibitory and additive kernels on the foveal side and scaled both sides to maintain a smooth contour (Figure 6A). The Gaussians underlying the inhibitory and additive kernels were identical in shape.

The responses of neurons were calculated similarly to the previously described model (*44*) using the same notation:

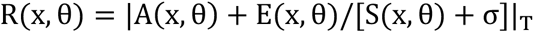

Where R is the response of a neuron, x represents its receptive field center, and θ its orientation preference. E is the excitatory drive, S is the suppressive drive, σ is a constant that determines the contrast gain, and T is the rectification threshold. A is the additive drive defined as:

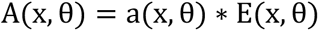

Where a(x, θ) is the asymmetrical additive kernel and * is convolution. Additional model parameters are presented below:

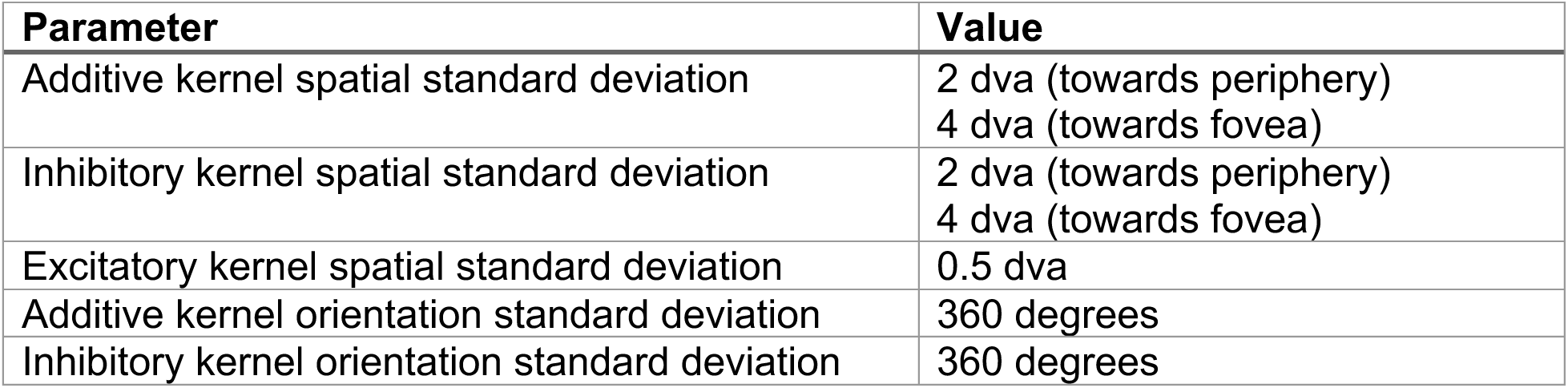
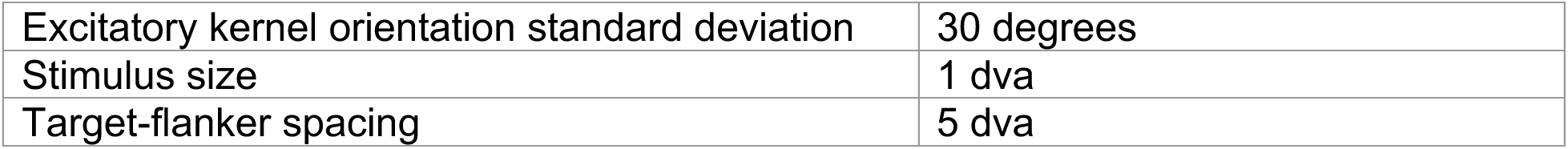

| Parameter | Value |
| --- | --- |
| Additive kernel spatial standard deviation | 2 dva (towards periphery)<br>4 dva (towards fovea) |
| Inhibitory kernel spatial standard deviation | 2 dva (towards periphery)<br>4 dva (towards fovea) |
| Excitatory kernel spatial standard deviation | 0.5 dva |
| Additive kernel orientation standard deviation | 360 degrees |
| Inhibitory kernel orientation standard deviation | 360 degrees |
| Excitatory kernel orientation standard deviation | 30 degrees |
| Stimulus size | 1 dva |
| Target-flanker spacing | 5 dva |

### Flanker-only firing responses

We calculated the firing rate of neurons from spikes occurring 0-300ms after flanker only stimulus onset. Firing rates were calculated for each stimulus position regardless of orientation. Neurons were split by cell class and layer and their firing rate was compared across stimulus locations.

